# Modulation of Pore Opening of Eukaryotic Sodium Channels by π-helices in S6

**DOI:** 10.1101/2023.03.25.534196

**Authors:** Koushik Choudhury, Lucie Delemotte

## Abstract

Voltage-gated sodium channels are heterotetrameric sodium selective ion channels that play a central role in electrical signaling in excitable cells. With recent advances in structural biology, structures of eukaryotic sodium channels have been captured in several distinct conformations corresponding to different functional states. The secondary structure of the pore lining S6 helices of subunit DI, DII, and DIV has been captured with both short π-helix stretches and in fully α-helical conformations. The relevance of these secondary structure elements for pore gating is not yet understood. Here, we propose that a π helix in at least DI-S6, DIII-S6, and DIV-S6 results in a fully conductive state. On the other hand, the absence of π-helix in either DI-S6 or DIV-S6 yields a sub-conductance state, and its absence from both DI-S6 and DIV-S6 yields a non-conducting state. This work highlights the impact of the presence of a π-helix in the different S6 helices of an expanded pore on pore conductance, thus opening new doors towards reconstructing the entire conformational landscape along the functional cycle of Nav Channels and paving the way to the design of state-dependent modulators.

**Graphical Abstract:** 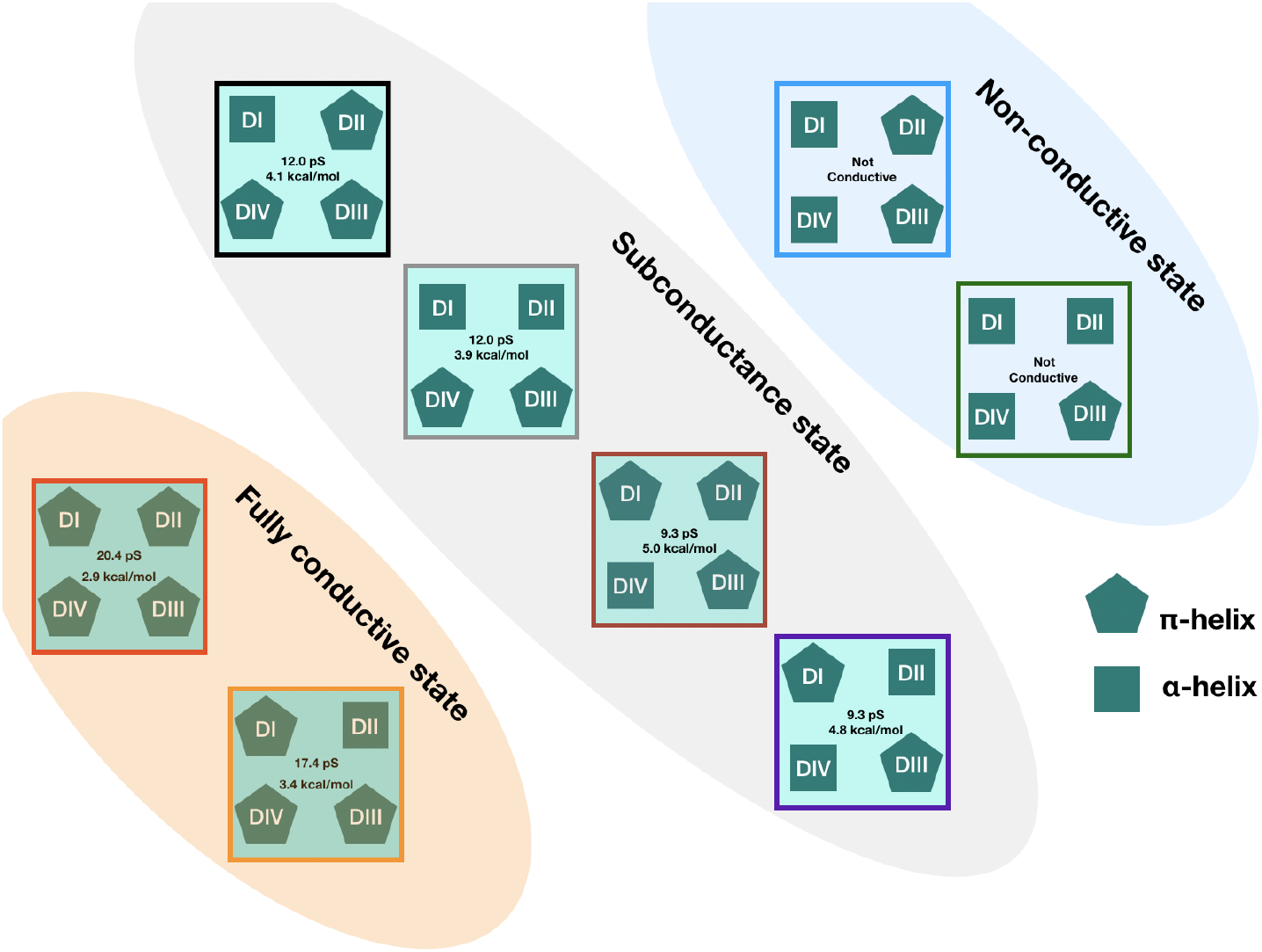

---

Voltage-gated sodium (Nav) channels are heterotetrameric membrane proteins that play an essential role in nerve impulse conduction in excitable cells (1). They selectively transport sodium ions across the membrane in response to membrane depolarization. The Nav channel family has nine isoforms (Nav1.1-Nav1.9) expressed in different excitable cells, whose sequence, structure, and function are highly conserved (2). Nav channels comprise ∼2000 residues arranged in a tetrameric architecture (Figure 1A) (3-19). Each subunit of Nav channels predominantly consists of two main domains (Figure 1B,C) - the voltage sensing domain (VSD) and the pore domain (PD). Subunit IV (DIV) contains the inactivation motif (IFM particle) binding site located on the DIII-DIV linker. The VSDs of each subunit consist of four helices labeled S1-S4 (Figure 1A,B,C), of which the S4 helix contains several Arginine and Lysine residues arranged every third position. These residues, so-called gating charges, sense changes in transmembrane voltage. The pore domain comprises the tetrameric assembly of two helices labeled S5 and S6 (Figure 1A,B,C). It can be divided into three main regions - selectivity filter, central cavity, and activation gate (Figure 1D). The VSD is connected to the pore domain through the S4-S5 linker (Figure 1A,B,C). Additionally, each subunit is connected to the next subunit through a disordered intracellular linker. The DIII-DIV linker features three hydrophobic residues (Isoleucine, Phenylalanine, and Methionine), which together form the IFM particle whose docking in its binding site appears to stabilize the inactivated state of the channel (Figure 1C) (3,5,6,8-10,12-19).

**Figure 1:**
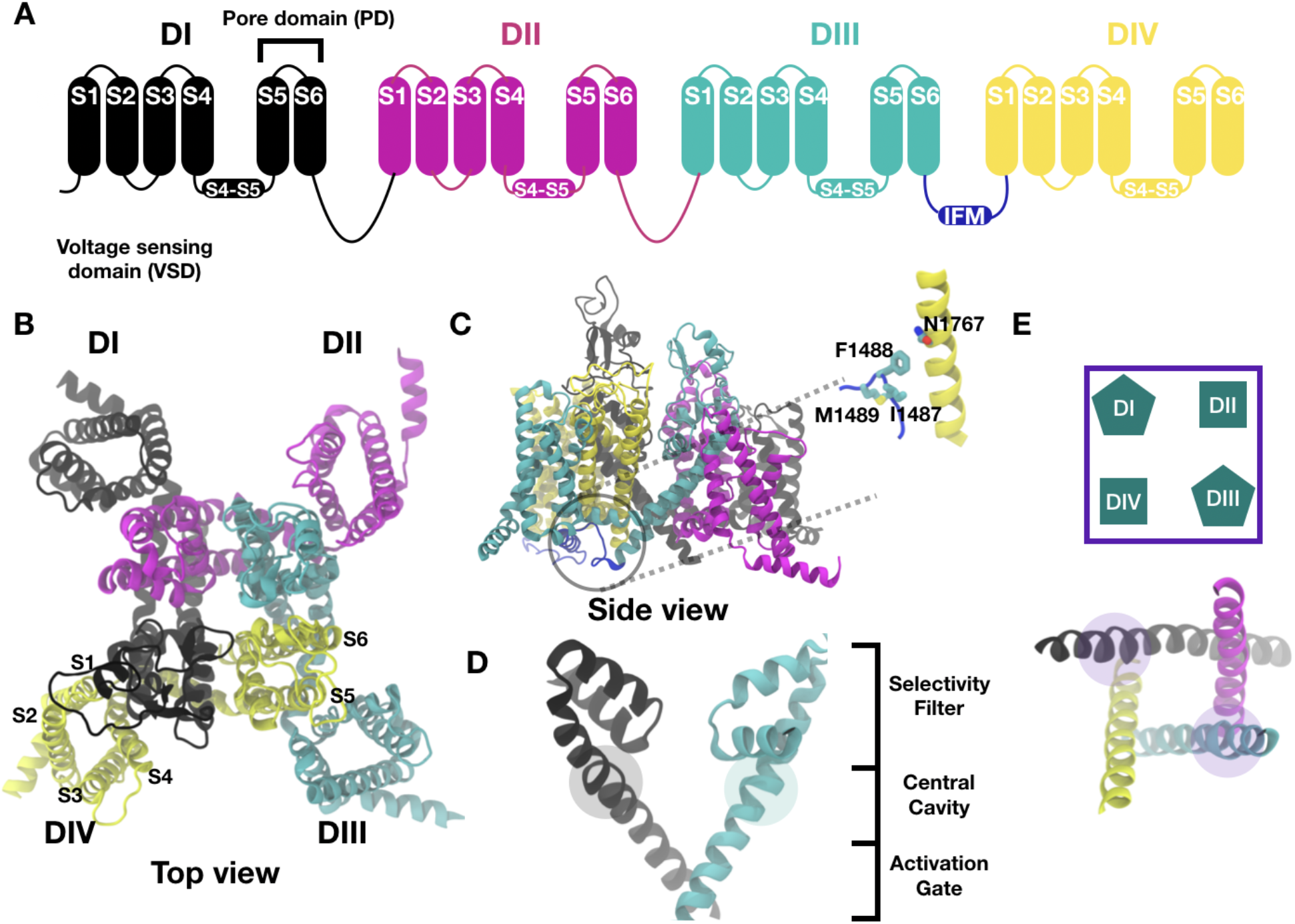
Architecture of eukaryotic Nav channels. **A**. Eukaryotic Nav channels exist as a single polypeptide chain arranged as a tetramer. **B, C**. Top (B) and Side (C) view of the experimentally resolved structure of cardiac sodium channel (Nav1.5, PDB - 7FBS). Inset shows a zoomed-in view of the bound inactivation particle (IFM) interacting with the conserved N1767 residue in subunit IV **D**. The pore lined by S6 helices can be divided into three main regions - selectivity filter, central cavity, and activation gate **E**. Top view of the pore of the experimental open state structure. The cartoon representation shows a simplified version of the structure. This representation will be used throughout this article. Pentagons represent S6s containing π-helices and squares represent fully *α*-helical S6s.

In response to membrane depolarisation, the VSDs activate via the displacement of the S4 helix towards the extracellular side. The VSDs of the different subunits have different activation kinetics. The activation of the VSDs of DI-DIII is sufficient to open the pore and allow free sodium ion permeation (20,21). Subsequent activation of the VSD of DIV leads to the channel’s entry in a second open state characterized by a lower conductance than the first one (21,22). This conformation allows rapid binding of the DIII-DIV linker IFM particle to the pocket formed by the S4-S5 linker of DIII and S6 of DIV, leading to pore closure. This phenomenon is known as fast inactivation (3) (Figure 1C).

The structures of different isoforms of eukaryotic Nav channels have been resolved in several conformational states (3-19), namely assigned as fast inactivated states (3,5,6,8-10,13-17,19), a closed inactivated state (18) and an open state (11). These different structures feature a variety of pore conformations. The DI, DII, and DIV S6 have been captured both with a fully α-helical content and with a short π-helix stretch below the selectivity filter (3,17,18,19). The DIII S6 helix, on the other hand, has always been captured containing a short π-helix across the different conformations of Nav channels (3-19). In detail, most of the experimental structures feature a π-helix in DI S6 and DIII S6 (3,5,6,8-10,13-16,19). The structures of Nav channels from the american cockroach feature a π-helix in all four S6 (4,7,12). The structure of the electrical eel Nav channel features a short π-helix in DII, DIII and DIV S6 (3). More recent structures feature a short π-helix in either DI, DII and DIII S6 or DI, DIII, DIV S6 (3,17,18,19).

In our recent studies of bacterial Nav channels NavMs (23) and NavAb (24), we found that introducing a short π-helical stretch in S6 of the channel resulted in the rotation of the S6 C-terminus and in increased pore hydration, allowing sodium ion permeation. This mechanism is consistent with that described in some TRP channels (25,26). The presence of π-helices in the pore lining helices of different channels with similar architecture (27) is considered to be conserved and to play an important role in drug binding (19,28). Inspired by these observations, we hypothesized that eukaryotic Nav channel opening might involve a transition to π-helix in one or more of the DI/DII/DIV S6 helices. Additionally, given the heterogeneity of eukaryotic Nav channels and the stabilization of different secondary structures in their S6 helices, we hypothesized that an α/π-helix conformation in different S6 helices might give rise to different sub-conductance levels, resulting in different open states.

Recently, a structure of the cardiac sodium channel isoform (Nav1.5) was captured with a wide open activation gate and a short π-helix in DI-S6 and DIII-S6 (a state we will henceforth refer to as 2pi_d1_d3) (Figure 1D) (11). In this study, we used molecular dynamics (MD) simulations to characterize this experimental open state by calculating the pore hydration, ion conductance, and ion permeation free energy. In addition, we investigated the same properties in several different models containing different numbers and combinations of π-helices in an attempt to assign potential open states. A general scheme is used to name these models wherein the first part of the name specifies the number of π-helices and the subsequent parts specify where on the S6 helices they are located. For example, 3pi_d2_d3_d4 refers to a model in which there are three π-helices, and in which these π-helices are located in DII, DIII, and DIV S6s.

To characterize the conductive properties of the Nav1.5 open state resolved experimentally (2pi_d1_d3, Figure 1E), we carried out 100 ns classical all-atom MD simulations and calculated the time-averaged water density of this model. Consistently with previous work, Ca positions were restrained to allow sidechain reorganization but prevent pore collapse (11). The water density profile along the pore axis shows that the pore is continuously hydrated. Although the water density near the activation gate drops to a value below that of bulk water density (Figure 2B, S1B), it is substantially higher than that in models of the open state of bacterial channels obtained experimentally (23,24,29), leading us to hypothesize that the presence of π-helices correlates with an increased water density. Indeed, the experimentally resolved open-state model of Nav1.5 featured two π-helix in DI-S6 and DIII-S6. To test this hypothesis, we tested the effect of removing the π-helix in DI-S6 using homology modeling (see Methods), leaving a model featuring a π-helix only in DIII-S6, namely 1pi_d3 (Figure 2A). Molecular dynamics simulations of this model revealed that this perturbation resulted in complete dehydration of the pore around the activation gate (Figure 2B, S1A). This could be attributed to the presence of hydrophobic residues facing the gate, namely a tetrad at the level of I409/F937/I1468/V1766 and another one helical turn below at V413/L941/I1472/I1770 (first panel (green), Figure 2C). The comparison between the 2pi_d1_d3 and the 1pi_d3 models suggested that a π-helix in DI-S6 is important for pore hydration. Introducing a π-helix in any S6 helix causes the rotation of the helix that follows the π-helix leading to a reorientation of the residues in that region. This also causes the rotation of the highly conserved Asparagine (23) towards the pore in a π-helical S6. A π-helix in DI-S6 causes the L410 and A414 to face the pore, while I409 and V413 face the pore when DI-S6 is *α*-helical (second panel (purple), Figure 2C). The Ala in position 414 thus reduces the hydrophobicity of the region given its smaller sidechain relative to V413 (second panel (purple), Figure 2C). Such a result prompted us to test the effect of inserting a π-helix in DII-S6 and DIV-S6. We thus created a 2pi_d2_d3 and a 2pi_d3_d4 model (Figure 2A). Molecular dynamics simulations of these revealed that the 2pi_d2_d3 pore was substantially dehydrated around the activation gate, yielding a hydration level comparable to the 1pi_d3 model. This suggested that a π-helix in DII-S6 is not essential for pore hydration (Figure 2B, S1C). The reason for this could be that a π-helix in DII-S6 rotates the I938 and I942 towards the pore, while F937 and I941 face the pore in the *α*-helical DII-S6 (fourth panel (blue), Figure 2C), leading to a minor change in hydrophobicity in the region. In contrast, in the 2pi_d3_d4 model, the pore was hydrated relative to the 2pi_d1_d3, 2pi_d2_d3 and 1pi_d3 model, suggesting that a π-helix in DIV-S6 is more conducive to pore hydration/ion permeation, when compared to π-helices inserted in DI-S6 and DII-S6 (Figure 2B, S1D). Consistently with our analysis, a π-helix in DIV-S6 causes the N1767 and A1771 to face the pore, to be compared to V1766 and I1770 facing the pore in *α*-helical DIV-S6 (third panel (grey), Figure 2C).

**Figure 2:**
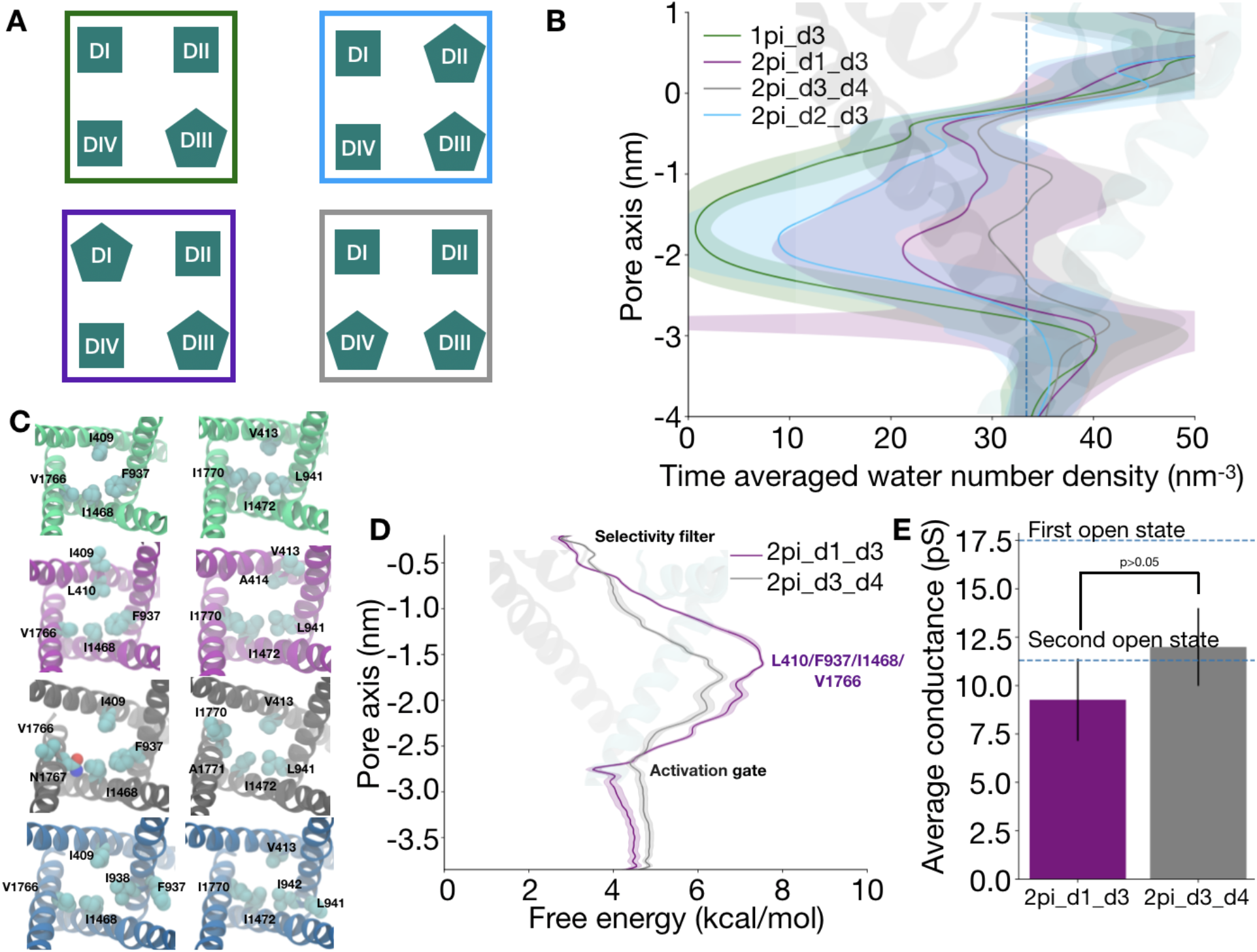
The presence of a π-helix in DI-S6 and DIV-S6 is important for ion conduction. **A**. Cartoon representation of the different pore conformations. **B**. Pore hydration around the activation gate in the different pore conformations **C**. Hydrophobic residues lining the pore in the 1pi_d3 model (green, 0 ns), 2pi_d1_d3 (purple, 100 ns), 2pi_d3_d4 (gray, 43 ns) and 2pi_d2_d3 (blue, 42 ns) **D**. Ion permeation free energy profiles in the models with a hydrated pore (2pi_d1_d3 in purple and 2pi_d3_d4 in gray). **E**. Average conductance values in the 2pi_d1_d3 (purple) and 2pi_d3_d4 (gray) from simulations. The blue dashed lines show the experimental conductance values for the first and second open state (21). Error bars represent the standard error across six replicates. p value < 0.05 indicates that there is a significant difference in the means between the two different models while a p value > 0.05 indicates that there is no significant difference.

To further investigate ion conduction properties of different models, we calculated ion permeation free energy using well-tempered metadynamics. Since the pore was dehydrated in the 2pi_d2_d3 model and 1pi_d3 model we assumed that ion conductance would be impeded by a high free energy barrier in the activation gate region. Indeed, our free energy profiles revealed free energy barriers of around 4.8 kcal/mol and 4.1 kcal/mol close to the activation gate in the 2pi_d1_d3 model and 2pi_d3_d4 model, respectively (Figure 2D, S10A, S10B). To further probe the propensity of these models to permeate ions and characterize the conductance level associated with such free energy barriers, we estimated the conductance directly by monitoring the number of ion permeation events over time. A previous MD simulation study of a bacterial Nav channel has indeed shown a linear relationship between current and voltage (30). We thus applied an electric field corresponding to a transmembrane potential difference of -500 mV and recorded ion permeation events. We observed around 1-5 ion permeation event across six 100 ns replicas in the 2pi_d1_d3 (Figure S3) and the 2pi_d3_d4 (Figure S4) model, resulting in conductance estimates of 9.3 +/- 2.1 pS and 10.5 +/- 2.0 pS respectively (Figure 2E). These conductance values are approximately two-thirds of the experimental conductance value of 17.5 pS (21), corresponding to 6.125 ion permeation events in 100 ns at -500 mV. A study of Nav1.4 channels proposed that pore gating involved three different open pore conformations (labeled as O, S1, and S2) with similar pore radius profiles but with conductance values decreasing from O to S2 to S1 (21). O and S2 open states are part of the main activation pathway, wherein O corresponds to the first open state and S2 corresponds to the second open state. The S1 open state, on the other hand, is not a part of the main activation pathway. Based on these observations and the fact that the 2pi_d1_d3 model has been captured experimentally, we propose that that structure corresponds to the S2 state. The 2pi_d3_d4 model is an alternative candidate for this state.

We hypothesized that the increased pore hydration in the 2pi_d1_d3 model relative to the bacterial experimental open models is due to the presence of a π-helix in DI-S6 and DIII-S6. Based on this, we sought to test how pore hydration and ion permeation are affected upon extending the presence of π-helices to both DII-S6 and DIV-S6. Inserting a π-helix in all four S6 helices (4pi, Figure 3A) resulted in an increase in pore hydration (Figure 3B, S2A), an increase in conductance (20.1 +/- 2.6 pS, Figure 3C, S5) and a decrease in the free energy barrier to ion permeation (2.9 kcal/mol, Figure 3D, S10C), relative to the 2pi_d1_d3 state. The conductance of this model is comparable to the experimentally measured conductance for the first open, O, state (Figure 3C).

**Figure 3:**
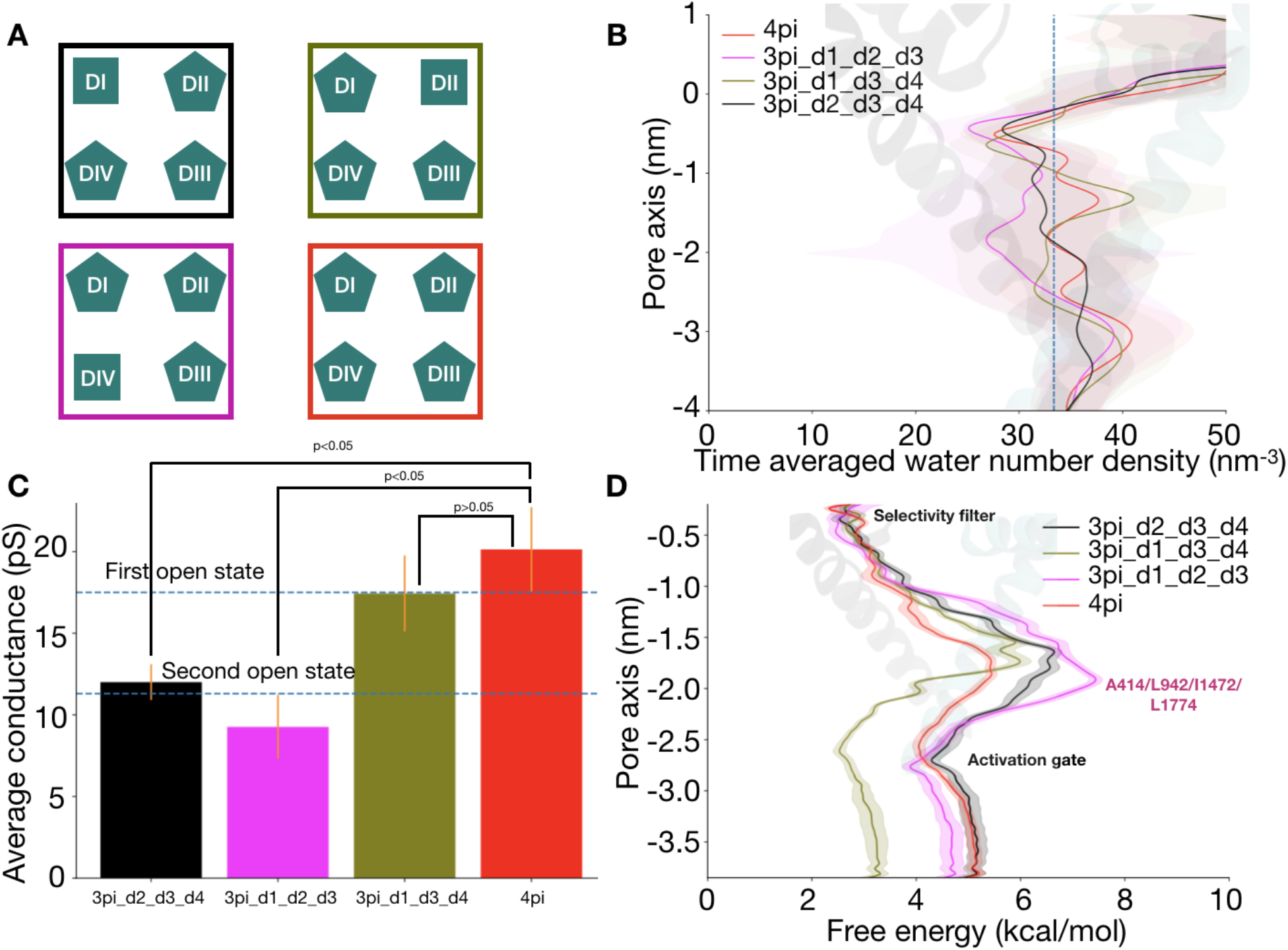
The presence of a π-helix in at least DI-S6, DIII-S6 and DIV-S6 are sufficient to generate a fully conductive pore. **A**. Cartoon representation of the different pore conformations **B**. Pore hydration around the activation gate in the different pore conformations (Black: 3pi_d2_d3_d4, Brown: 3pi_d1_d2_d3, Orange: 3pi_d1_d3_d4, Red: 4pi) **C**. Average conductance values in the different models across six replicates from simulations. The blue dashed lines show the experimental conductance values for the first and second open state (21). Error bars represent the standard error across six replicates. p value < 0.05 indicates that there is a significant difference in the means between the two different models while a p value > 0.05 indicates that there is no significant difference. **D**. Ion permeation free energy profile in the different models

To investigate the contribution of π-helices in each S6 to pore hydration and ion conduction properties, we tested different models wherein the π-helix is removed from a single S6 helices, resulting in models with π-helices in three of four S6. Upon removing the π-helix either from DIV-S6 (3pi_d1_d2_d3) or DI-S6 (3pi_d2_d3_d4), pore hydration and ion conductance dropped, accompanied by an increase in the free energy barrier for ion permeation. This suggested that the presence of a π-helix in DIV-S6 and DI-S6 might be essential to model an open state. The pore hydration (Figure 3B, S2C), ion conductance (9.3 +/- 2.0 pS, Figure 3C, S6 versus 12.0 +/- 1.1 pS, Figure 3C, S7) and free energy barrier for ion permeation (5 kcal/mol vs 4.1 kcal/mol, Figure 3D, S10D) are comparable in the 3pi_d1_d2_d3 and the 3pi_d2_d3_d4 models. Comparing the 2pi_d2_d3 model to the 3pi_d2_d3_d4 model suggests that a π-helix in DIV-S6 could be important (Figure 4). Additionally, comparing the 2pi_d2_d3 model to the 3pi_d1_d2_d3 model suggests that a π-helix in DI-S6 might be important for conduction as removing the π-helix in DI-S6 from 3pi_d1_d2_d3 model resulted in pore dehydration (Figure 4).

**Figure 4:**
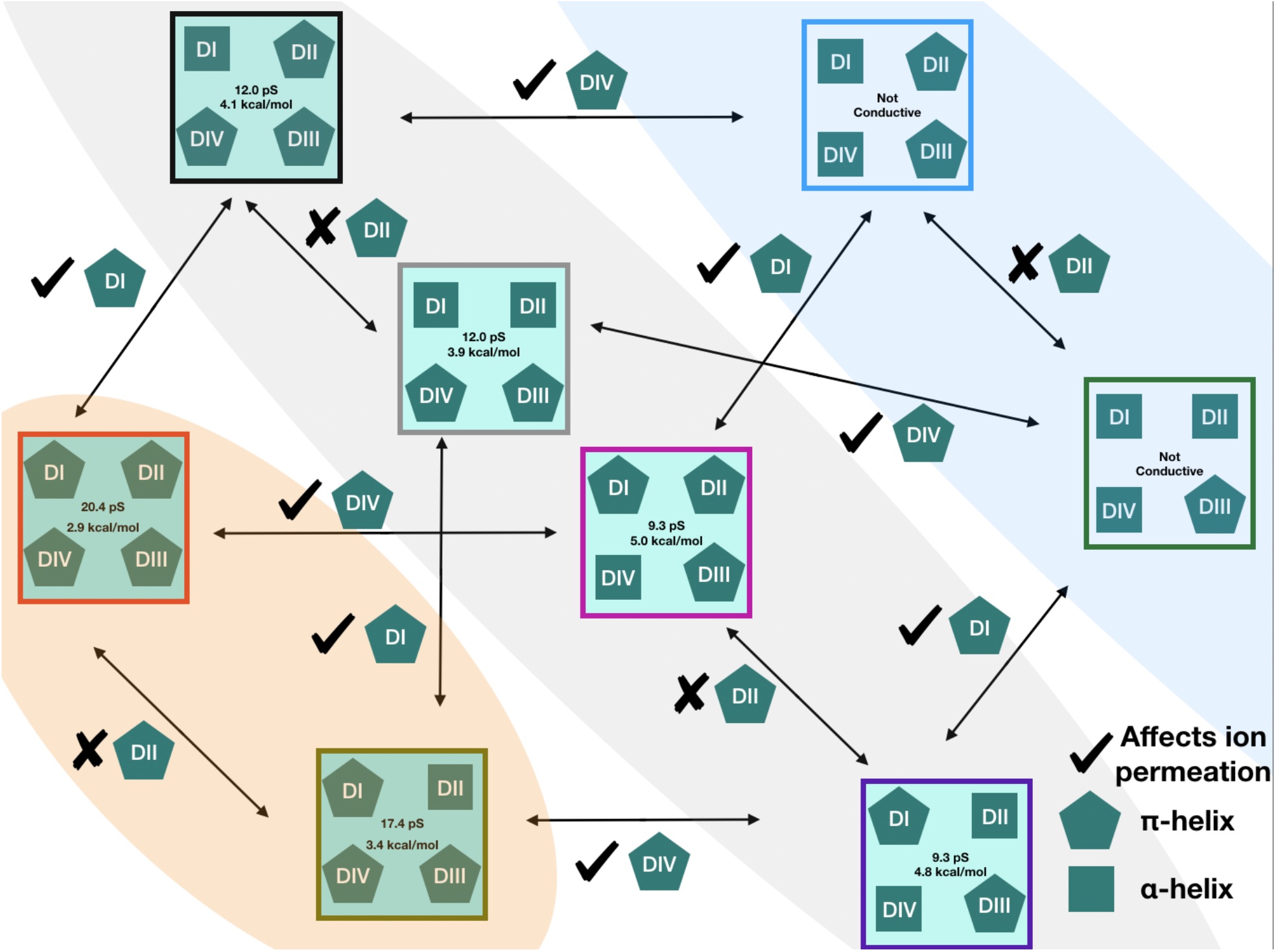
Flowchart showing the effect of removing/adding π-helices to different S6 helices. The orange-shaded region shows the pore conformations that have a fully-conductive pore. The blue-shaded region shows the non-conductive pore conformations. The gray-shaded region shows pore conformations that are sub conductive. The pentagons represent π-helices and the squares represent *α*-helices in S6 helix. The pentagon following a tick mark shows that a π-helix in the corresponding S6 can affect ion permeation while a cross mark signifies that it does not affect ion permeation.

On the other hand, removing the π-helix from DII-S6 from the 4pi model (resulting in 3pi_d1_d3_d4) leads to pore hydration (Figure 3B, S2B), ion conductance (17.4 +/- 2.3 pS, Figure S8) and free energy barrier for ion permeation (3.4 kcal/mol, Figure 3D,S10F) that are not substantially affected, suggesting that a π-helix in DII-S6 might not be essential for conductive properties. Additionally, pore hydration, conductance, and barrier for ion permeation remained relatively unaffected between the 3pi_d1_d2_d3 model and 2pi_d1_d3 model, and between the 3pi_d2_d3_d4 model and 2pi_d3_d4 model (Figure 4). These results further support our claim that a π-helix in DII-S6 is not essential for ion permeation.

In the recent structures of Nav1.7 featuring a π-helix in DIV-S6, the π-helix is shifted downwards by a helical turn relative to the π-helix in the 3pi_d1_d3_d4 model (17). Indeed, the π-helix is localized at N1767, in contrast to being localized at F1762 in the 3pi_d1_d3_d4 model (Figure S11A). We thus tested the effect of shifting the π-helix a helical turn downward in the 3pi_d4 model, placing it at the level of N1767. We observed no effect on pore hydration (Figure S11B), ion permeation free energy, and conductance (Figure S9, S11C,S11D). The precise localization of the π-helix along the helix thus does not appear to affect the propensity of the pore for ion conduction.

We have built possible models of Nav1.5 open-pore states, in addition to the experimentally-determined model containing two π-helices in DI and DIII. The various models have different S6 secondary structure compositions, resulting in properties consistent with non-conductive, sub-conductive, or fully-conductive states. One similarity among the different sub-conductive states (Figure 4, grey shade) is that the pore domain features a π-helix in either the DI-S6 or DIV-S6. The similarity between the two non-conductive (Figure 4, blue shade) states is that they do not contain a π-helix in both DI-S6 and DIV-S6. For a fully-conductive state (Figure 4, orange shade) a π-helix in both DI-S6 and DIV-S6 is necessary. Additionally, we also conclude that the conductance of the pore is likely directly related to the number of π-helices in the S6s of different subunits. Although the simulations in this study were performed under restraints, the results presented in this study reveal an important role of the π-helix on pore hydration/ion permeation given an expanded pore conformation. Starting from either the 4pi model or the 3pi_d1_d3_d4 model, a π- to *α*-helix transition in DIV-S6 only is enough to increase the barrier for ion permeation sufficiently to lower the conductance value. For fast inactivation to occur, the second open state must be attained. Here, we surmise that a π to *α*-helix transition in DIV-S6 reduces the conductance, corresponding to the transition from first to second open state. Thus, a π to *α*-helix transition in DIV-S6 might be an important step towards fast inactivation, in addition to VSD-DIV activation. We thus suggest that a π-helix in DI-S6, DIII-S6 and DIV-S6 with an expanded pore is enough to allow sodium ion permeation and is likely to correspond to the first open state. Activation of VSD-DIV and π to *α*-helix transition in DIV-S6 while maintaining an expanded pore will then lead to the sub-conductance state named the second open state. This conformational change will possibly allow IFM binding and hence lead to fast inactivation.

## Supporting Information

Experimental methods, time series plots of water number density and ion permeation, convergence profiles of the free energies, and additional comparisons of time-averaged pore hydration, average conductance, and ion permeation free energy profile turn

## Acknowledgements

We acknowledge SciLifeLab and the Swedish Research Council to LD (VR 2018-04905, VR 2019-02433 and VR 2022-04305) for funding. MD simulations were performed on resources provided by the Swedish National Infrastructure for Computing (SNIC) at PDC Center for High Performance Computing (PDC-HPC). We also acknowledge PRACE for awarding us access to Piz Daint at CSCS, Switzerland.

## Author contributions

K.C and L.D designed the research. K.C performed the simulations and analyzed the results. K.C and L.D interpreted the data and wrote the manuscript.

## Competing interests

The authors declare to have no competing interests.

## Notes

### Competing Interest Statement

The authors have declared no competing interest.

### Summary of Updates

Revised after peer-review.

