## Supplementary figures and Methods for "Modulation of Pore Opening of Eukaryotic Sodium Channels by π-helices in S6"

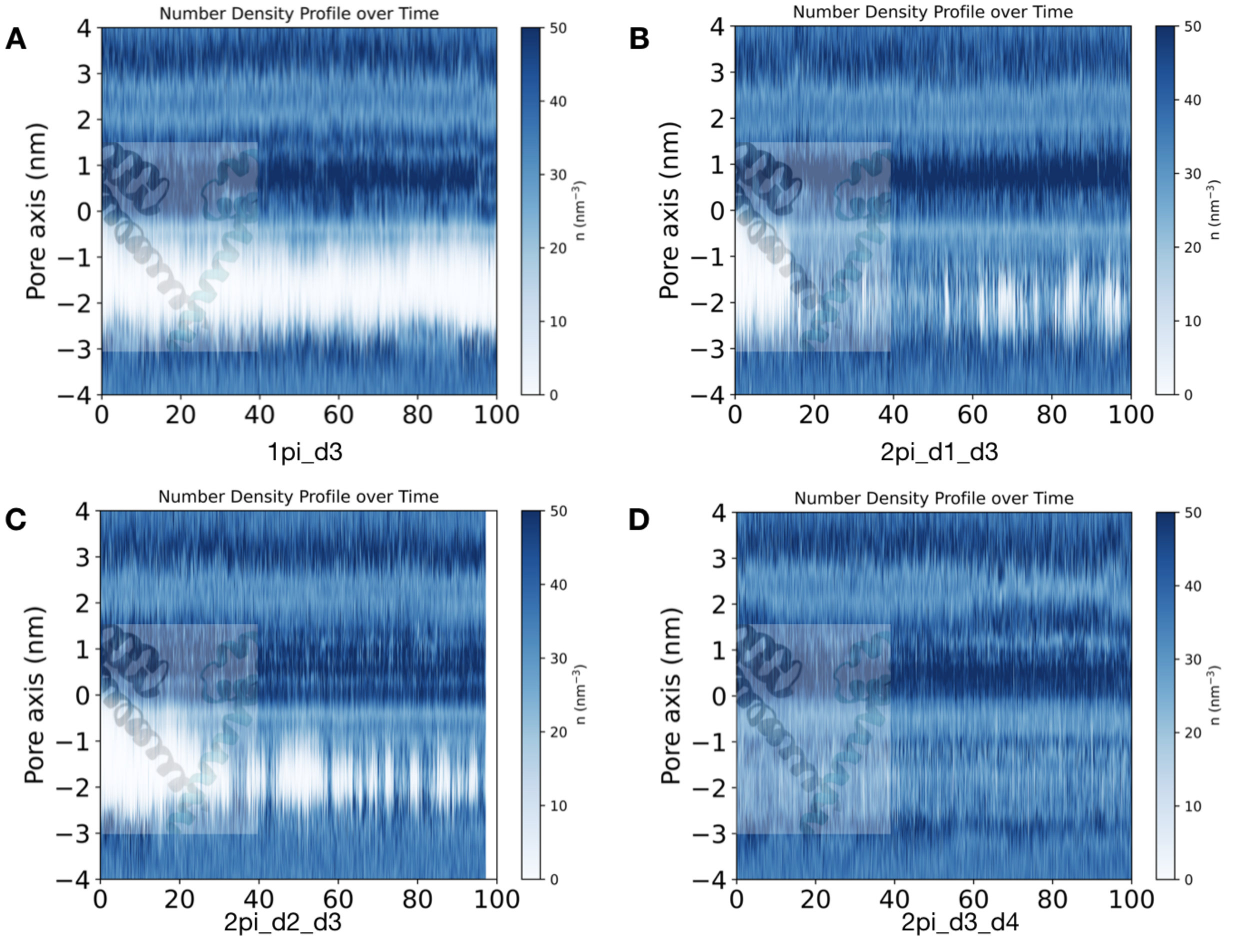


**Figure S1:** Pore hydration as a function of time in the **A.** 1pi_d3 model, **B.** 2pi_d1_d3 model, **C.** 2pi_d2_d3 model, **D.** 2pi_d3_d4 model

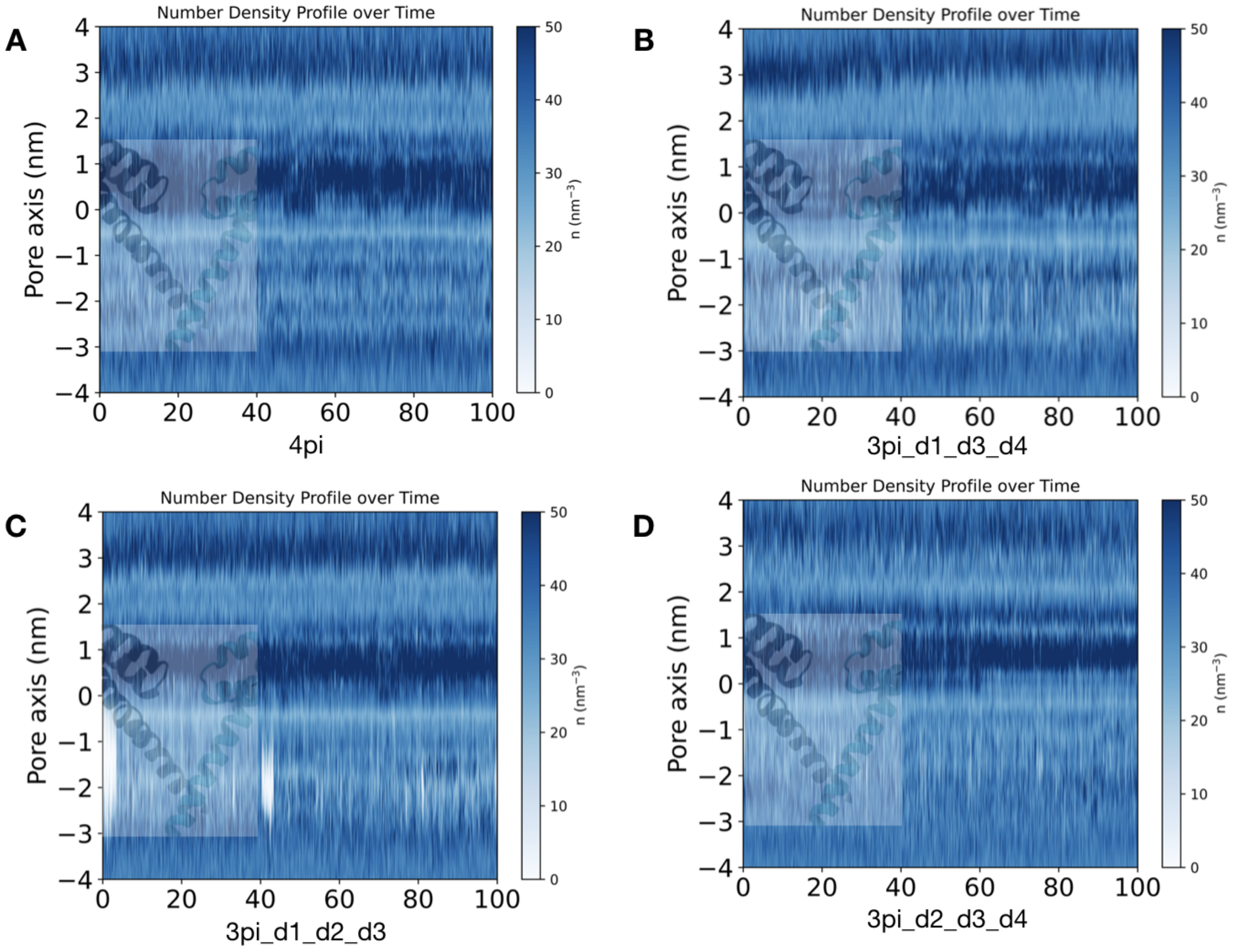


**Figure S2:** Pore hydration as a function of time in the **A.** 4pi model, **B.** 3pi_d1_d3_d4 model, **C.** 3pi_d1_d2_d3 model, **D.** 3pi_d2_d3_d4 model

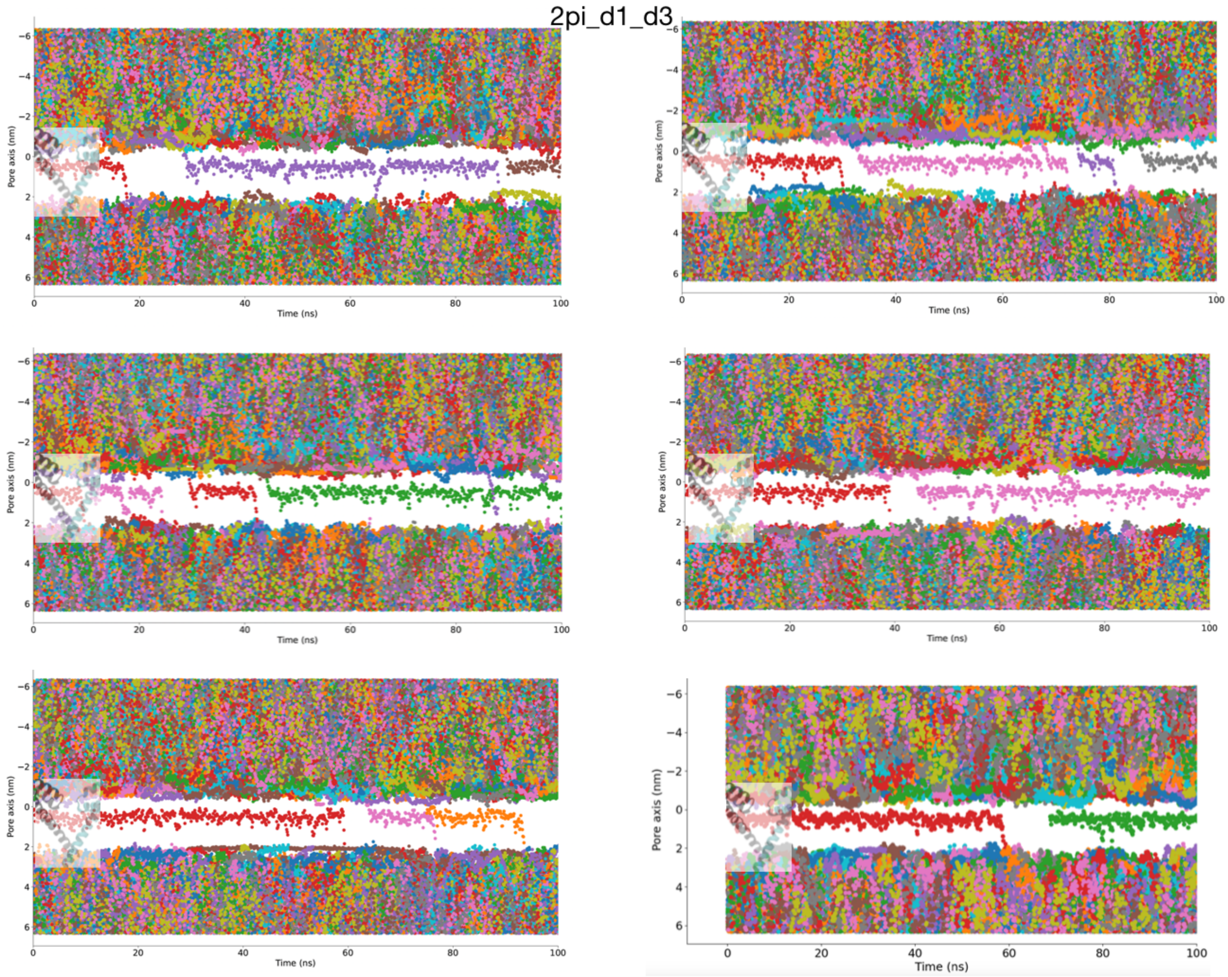


**Figure S3:** Sodium ion permeation trajectories in the different 100 ns long simulations replicates of 2pi_d1_d3 model at -500 mV

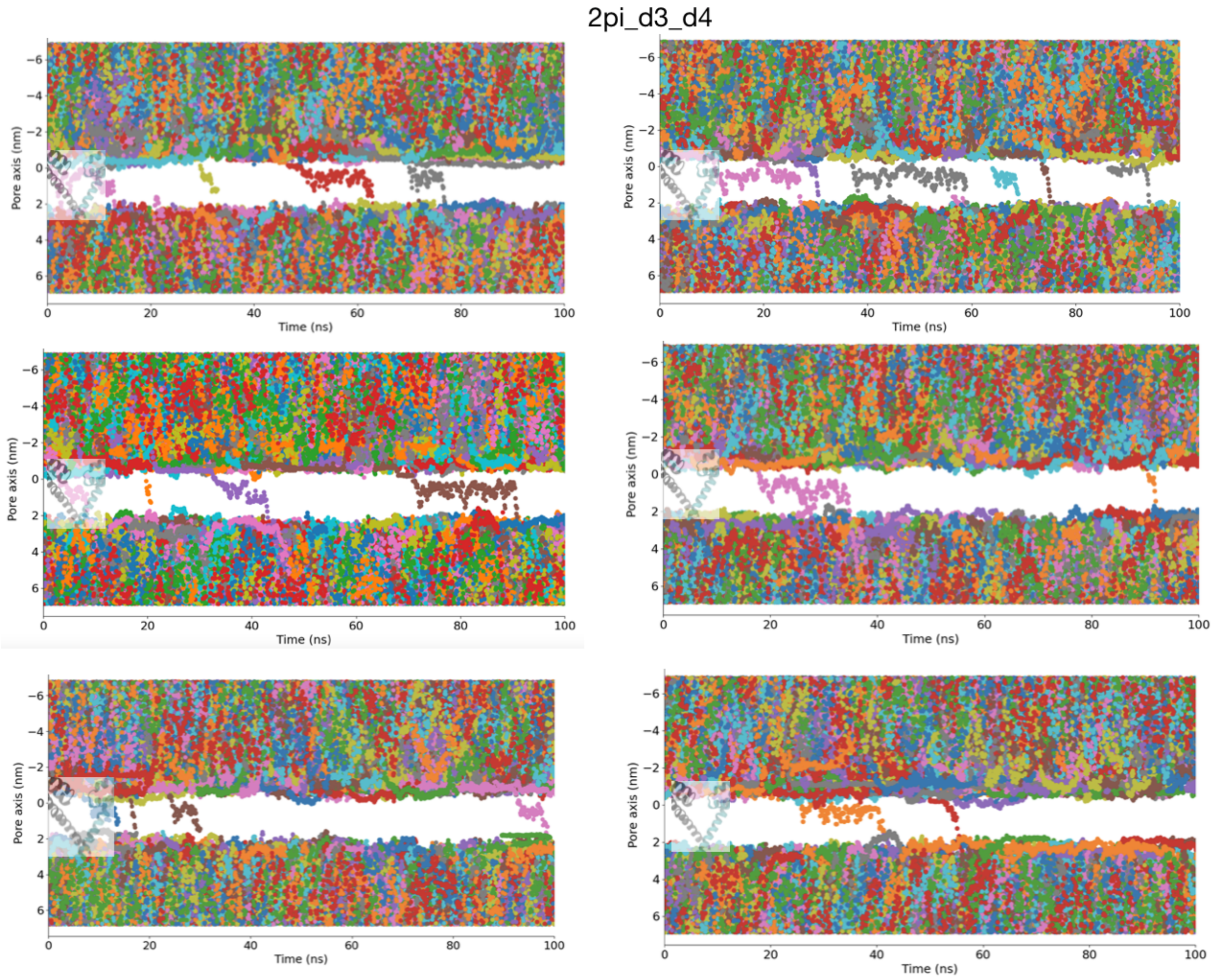


**Figure S4:** Sodium ion permeation trajectories in the different 100 ns long simulations replicates of 2pi_d3_d4 model at -500 mV

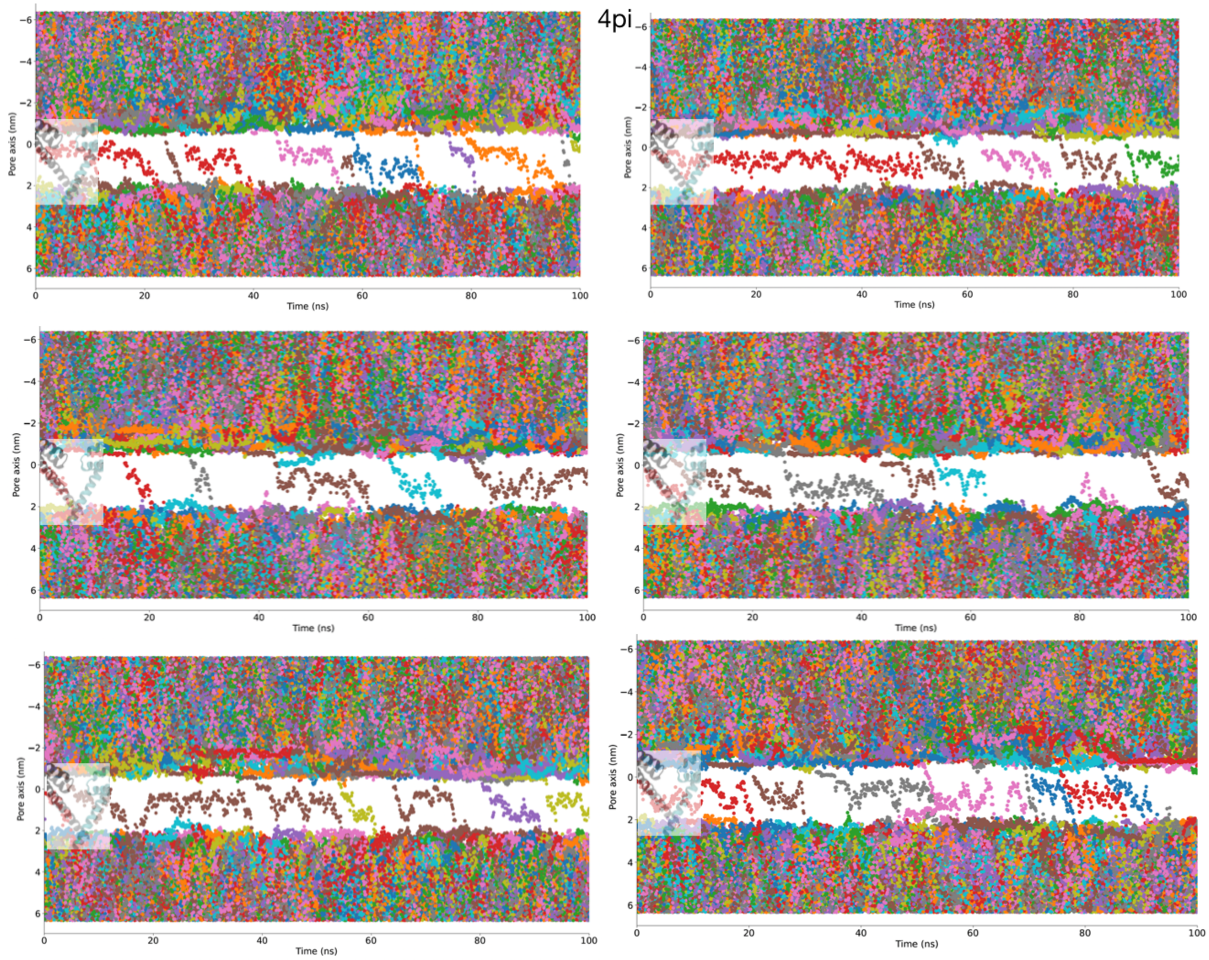


**Figure S5:** Sodium ion permeation trajectories in the different 100 ns long simulations replicates of 4pi model at -500 mV

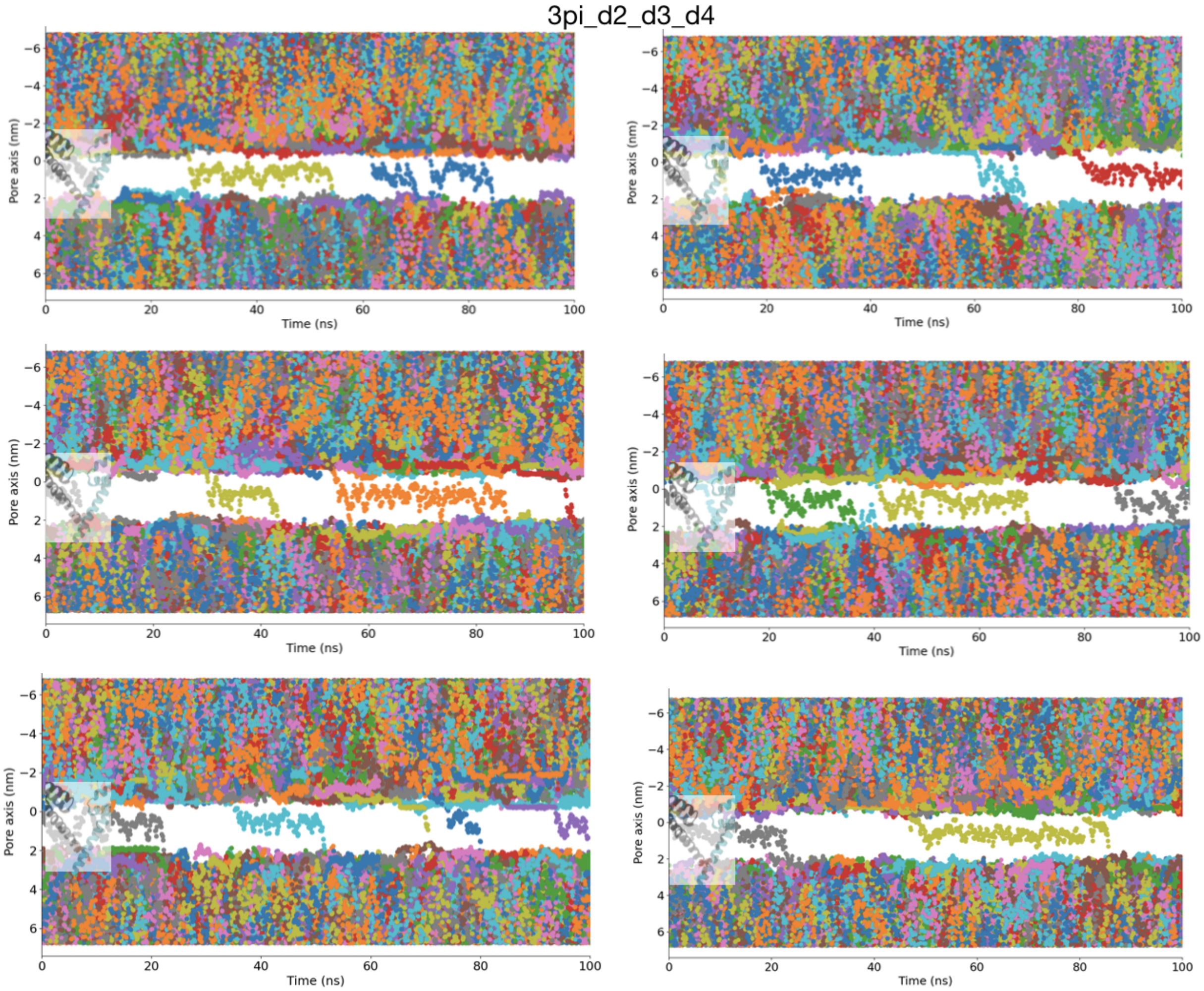


**Figure S6:** Sodium ion permeation trajectories in the different 100 ns long simulations replicates of the 3pi_d2_d3_d4 model at -500 mV

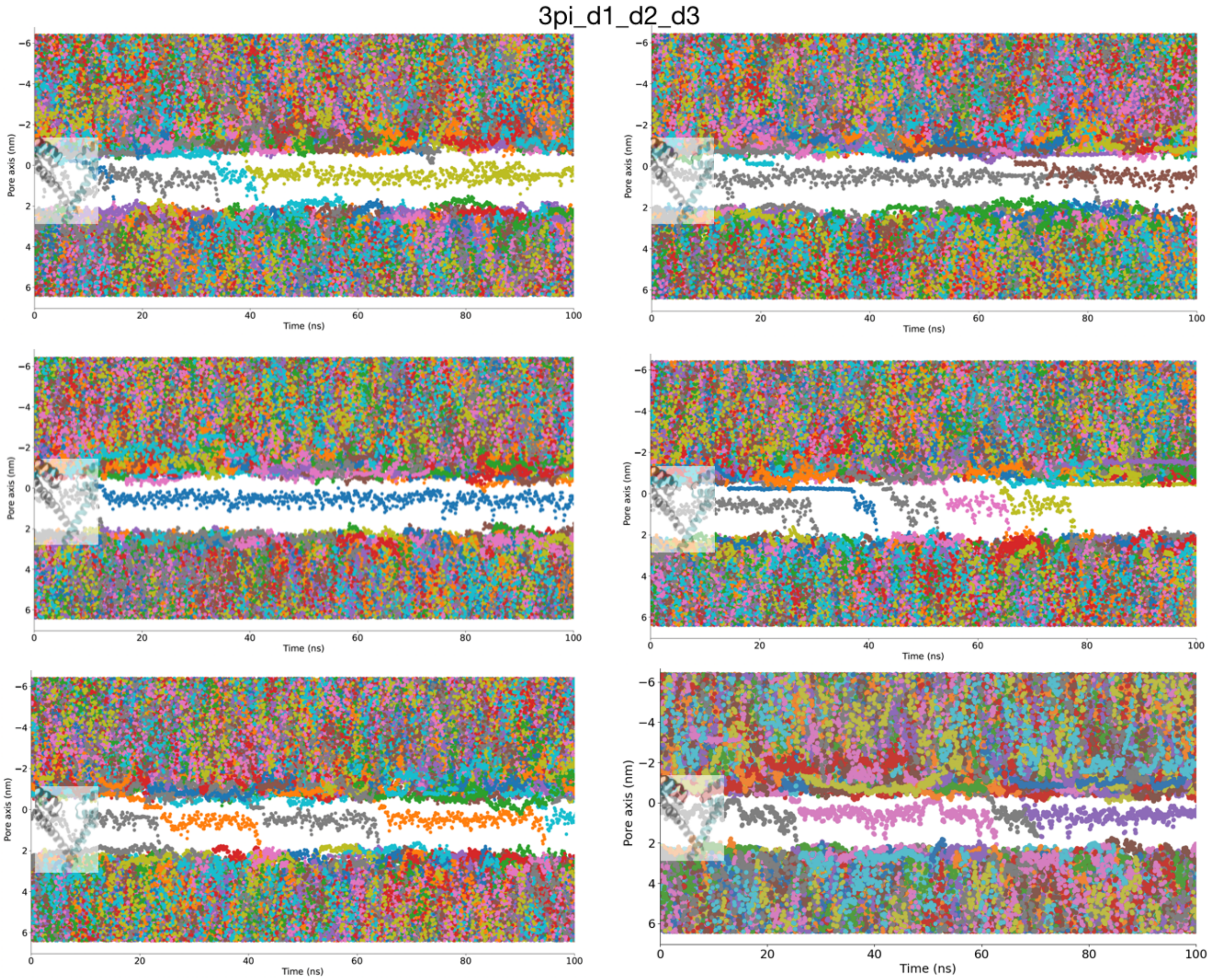


**Figure S7:** Sodium ion permeation trajectories in the different 100 ns long simulations replicates of the 3pi_d1_d2_d3 model at -500 mV

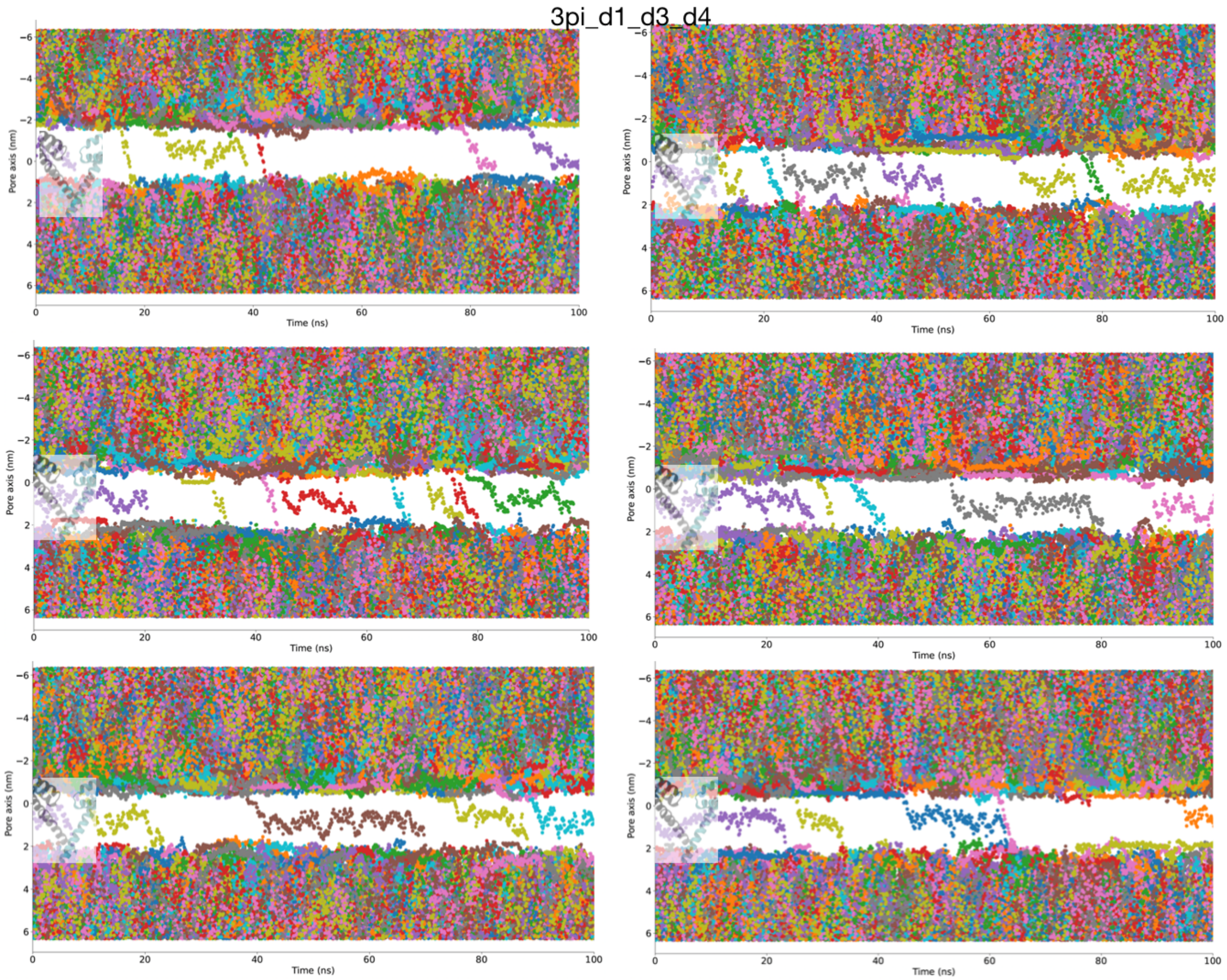


**Figure S8:** Sodium ion permeation trajectories in the different 100 ns long simulations replicates of the 3pi_d1_d3_d4 model at -500 mV

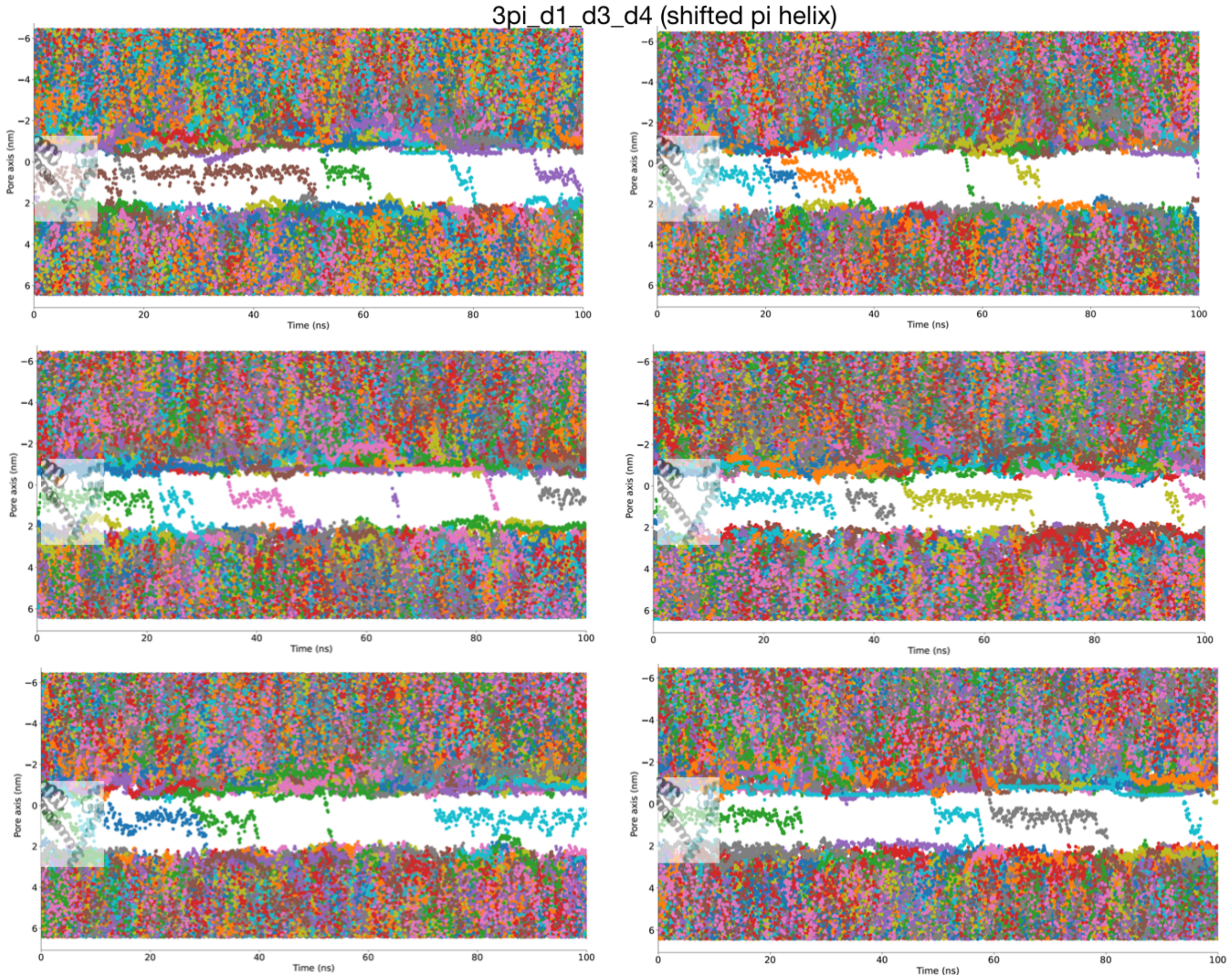


**Figure S9:** Sodium ion permeation trajectories in the different 100 ns long simulations replicates of the 3pi_d2_d3_d4 model in which the $\pi$-helix was shifted by a helical turn at -500 mV

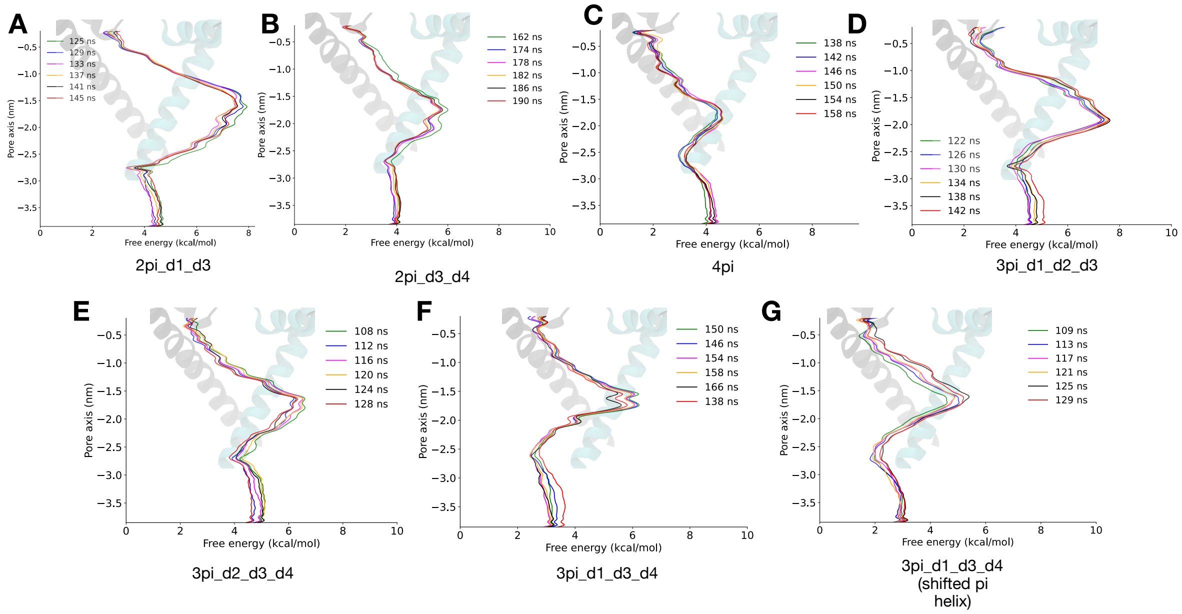


**Figure S10:** Convergence of the ion permeation free energy profiles in the different models **A.** 2pi_d3 **B.** 2pi_d3_d4 **C.** 4pi **D.** 3pi_d1_d2_d3 **E.** 3pi_d2_d3_d4 **F.** 3pi_d1_d3_d4 **G.** 3pi_d1_d3_d4 (shifted pi helix). The profiles were shifted to align to one another.



S1
0

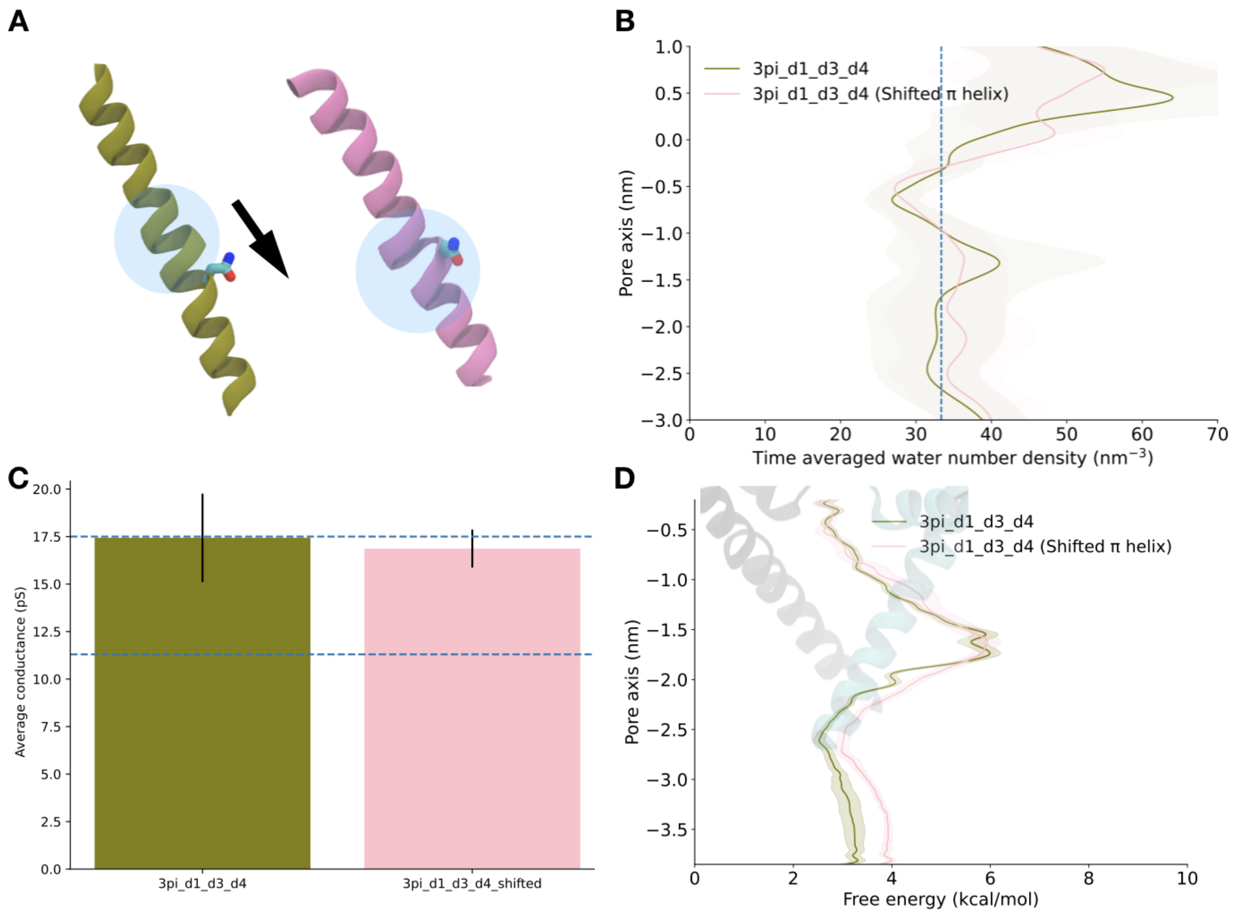


**Figure S11: A.** Difference between **B.** Pore hydration around the activation gate in the different pore conformations **C.** Average conductance values in the different models across six replicates. Error bars represent the standard error across six replicates. p value < 0.05 indicates that there is a significant difference in the means between the two different models while a p value > 0.05 indicates that there is no significant difference. **D.** Ion permeation free energy profile in the two models.

### Methods

#### Model building

The experimental open state model of Nav1.5 (PDB - 7FBS) was used in this study. Alternative open pore conformations were built by introducing $\pi$-helix in DII only, then in DIV only and finally in both DII and DIV. These modifications were introduced to the experimental open state model (7FBS). This resulted in four different models with different configurations of $\pi$-helix in S6 helices - $\pi$-helix in DI and DIII S6 (2pi_d1_d3 model), $\pi$-helix in DI, DII, DIII S6 (3pi_d1_d2_d3 model), $\pi$-helix in DI, DIII, DIV S6 (3pi_d1_d3_d4 model), $\pi$-helix in DI, DII, DIII and DIV S6 (4pi model) and $\pi$-helix in DI, DIII and $\pi$-helix at N1767 of DIV S6. This was done following a protocol implemented in previous work on NavMs using MODELLER 9.22 (1). To build models in which DI S6 was α-helical, a homology model of Nav1.5 was built using the Nav1.4 electric eel structure as template (PDB - 5XSY) using Swissmodel (2). The DI from this homology modeled structure was then used to substitute the DI in 4pi, 2pi_d2_d3, 3pi_d1_d3_d4 and 3pi_d1_d2_d3 models, resulting in 3pi_d2_d3_d4, 1pi_d3, 2pi_d3_d4 and 2pi_d2_d3 models respectively. All the models modeled using the Swissmodel structure are made available at https://osf.io/dx2hc/. The residues around the missing intracellular domains were capped. The residues forming the IFM particle were missing in the experimental open state model of Nav1.5. In the open state, the IFM particle is not supposed to be bound thus we decided not to build these missing residues and remove the DIII-DIV linker altogether. All the missing residues were built using CHARMM-GUI (3).

#### System preparation

Each model was embedded in a homogenous lipid bilayer consisting of 400 1-palmitoyl-2-oleoyl-*sn*-glycero-3-phosphocholine (POPC) using the CHARMM-GUI Membrane builder (3). Each system was hydrated by adding ~45 Å layers of water to either side of the membrane. Lastly, the systems were neutralized with 150 mM NaCl. The Charmm36 force field was used to describe interactions between protein (4), lipids (5) and ions. Non-bonded fixes (NBFix) were considered in the description of interactions of sodium ions with carboxyl and carbonyl groups (6). The TIP3P model was used to describe all water molecules (7).

**Simulation details**



For all the models, equilibration was carried out in six steps according to the default CHARMM-GUI protocol (3). During equilibration, a time step of 2 fs was used, pressure was maintained at 1 bar through Berendsen pressure coupling, temperature was maintained at 300 K through Berendsen temperature coupling (8) with the protein, membrane and solvent coupled and the LINCs algorithm (9) was used to constrain the bonds involving hydrogen atoms. For long range interactions, periodic boundary conditions and particle mesh Ewald (PME) were used (10). For short range interactions, a cut-off of 12 Å was used. Production simulations were run with backbone position restraints for different systems as shown in Table 1, using Parinello-Rahman pressure coupling (11) and Nose- Hoover temperature coupling (12). Simulations were performed using GROMACS 2020.5 and GROMACS 2021.5 (13,14). Some simulations were performed under a transmembrane voltage of -500 mV, via application of an external electric field to calculate conductance. The transmembrane voltage is calculated as the product of electric field and the simulation box length along which the field is applied.

**Table 1: Summary table of equilibrium simulations performed. Restrained production simulations were used to prevent pore collapse (15).**

| **System** | **Production** |
|---|---|
| 1pi_d3 model | 100 ns |
| 2pi_d1_d3 model | 100 ns |
| 2pi_d2_d3 model | 100 ns |
| 2pi_d3_d4 model | 100 ns |
| 3pi_d1_d2_d3 model | 100 ns |
| 3pi_d1_d3_d4 model | 100 ns |
| 3pi_d2_d3_d4 model | 100 ns |
| 4pi model | 100 ns |

| | |
|---|---|
| 3pi_d1_d3_d4 model (shifted $\pi$-helix) | 100 ns |
| 2pi_d1_d3 model (-500 mV) | 100 ns X 6 replicas |
| 2pi_d3_d4 model (-500 mV) | 100 ns X 6 replicas |
| 3pi_d1_d2_d3 model (-500 mV) | 100 ns X 6 replicas |
| 3pi_d1_d3_d4 model (-500 mV) | 100 ns X 6 replicas |
| 3pi_d2_d3_d4 model (-500 mV) | 100 ns X 6 replicas |
| 4pi model (-500 mV) | 100 ns X 6 replicas |
| 3pi_d1_d3_d4 model (shifted $\pi$-helix) (-500 mV) | 100 ns X 6 replicas |

**Ion permeation free energy calculations**

The ion permeation free energy along the pore axis was also calculated using multiple walker well-tempered metadynamics (16,17) in GROMACS 2020.5 patched with PLUMED 2.7.3 (18,19). Each bias acts on the z-axis defined using the center-of-mass *z*-distance between one sodium ion and C-α atoms of S1178 residues in NavAb. The hills height and width was set at 1.2 kJ/mol and 0.05 nm respectively. A bias factor of 15 was used. Upper and lower restraints (as shown in the corresponding x-axes of the free energy plots) were applied to restrict the sampling space. To keep the sodium ion close to the pore axis, the coordinate radial distance was restrained to stay below 6 Å by restraining the x and y component of center-of-mass distance. The free energy profile was estimated using the sum_hills tool in PLUMED. The average free energy profile was calculated by taking the data from the last 20 ns (per walker) in intervals of 4 ns and the error bars were estimated by calculating the standard deviation.

**Table 2: Summary table of equilibrium simulations performed. Restrained production simulations (where indicated) were used to prevent pore collapse.**

| System | Enhanced Sampling technique | Production |
|---|---|---|
| 2pi_d1_d3 model | Well-tempered metadynamics | 145 ns × 6 walkers |
| 3pi_d1_d2_d3 model | Well-tempered metadynamics | 142 ns × 6 walkers |
| 3pi_d1_d3_d4 model | Well-tempered metadynamics | 166 ns × 6 walkers |
| 4pi model | Well-tempered metadynamics | 158 ns × 6 walkers |
| 3pi_d1_d3_d4 model (shifted $\pi$-helix) | Well-tempered metadynamics | 129 ns × 6 walkers |
| 2pi_d3_d4 model | Well-tempered metadynamics | 190 ns × 6 walkers |
| 3pi_d2_d3_d4 model | Well-tempered metadynamics | 128 ns × 6 walkers |

**Analysis**

The water number density was calculated using the channel annotation package (CHAP) (20). CHAP was also used to classify S6 residues as pore-lining and pore-facing. For all the analyses, frames were extracted every 100 ps from the trajectories of the respective models. The number of ion permeation events was calculated by plotting the trajectory of sodium ions and then counting the number of sodium ions that traversed the entire pore length. Conductance was calculated using the following formula - $N \times 1.6 \times 10^{-19} / t \times 10^{-9} \times V$, where N is the number of ion permeation events, t is simulation time in seconds and V is the voltage in V. This gives the conductance in S which is then converted to pS. The average conductance was calculated across replicas and the error bars were estimated by calculating the standard error. A two-sample t-test was performed to analyze the statistical significance of the average conductance calculations.



S1
6
